# Thyroid hormone and ALK5 inhibitor improve maturation of human pluripotent stem cell derived hepatocytes

**DOI:** 10.1101/2022.04.28.489845

**Authors:** Sarah Withey, David Gerrard, Hannah Leeson, Rebecca Atkinson-Dell, Sean Harrison, Melissa Baxter, Ernst Wolvetang, Neil Hanley

## Abstract

Hepatocytes derived from human pluripotent stem cells (PSCs) hold great promise for modeling human liver disease, *in vitro* hepatotoxicity testing, and future cellular therapy. However, current protocols generate hepatocyte-like cells (HLCs) that resemble fetal hepatocytes, and thus do not accurately recapitulate the molecular identity and functions of the adult liver. To address this, we compared the transcriptomes of human fetal and adult liver to PSC-derived HLCs during progressive stages of *in vitro* differentiation. This revealed that during the final stages of *in vitro* differentiation the hepatic transcription factors HNF4A and CEBPA were sub-optimally expressed. Computational analyses predicted that ALK5i II (TGF-β receptor inhibitor) and thyroid hormone (T3) would be able to rectify this and improve HLC maturation. We next show that application of these molecules during hepatocyte differentiation indeed increases CEBPA and HNF4A mRNA and protein expression, and that these HLCs show enhanced albumin secretion, a 25-fold increase in CYP3A4 activity, and 10 to 100-fold increased expression of mature hepatic markers. We demonstrate that this improved maturation is effective across different cell lines and HLC differentiation protocols, and exemplifies that our approach provides a tractable template for identifying and targeting additional factors that that will fully mature human liver cells from human pluripotent stem cells.

## Introduction

Cellular models of hepatotoxicity need to faithfully capture the phenotype of hepatocytes. The ‘gold standard’ cell model for hepatotoxicity studies has been primary human hepatocytes, but their limited availability, inter-individual variability, and de-differentiation once in culture beyond a few days [1], has spurred the search for alternative models. The directed differentiation of human pluripotent stem cells (PSCs) towards hepatocyte-like cells (HLC) can overcome these drawbacks. We and others have shown that that progression of human pluripotent stem cells through the three main stages of early liver development: definitive endoderm differentiation, hepatocyte progenitor generation (with similarity to hepatoblasts), and hepatocyte specification [2-7], can be accomplished through stepwise addition of soluble growth factors, cytokines or small molecules. To enhance hepatocyte maturity dexamethasone (DEX), oncostatin M (OSM), HGF [3-5, 8-11] or microbial-derived substances have often been applied [12]. Despite these advances, we [13] and others [7] have found that differentiation protocols generate HLC with phenotypes that fall short of adult hepatocytes. Indeed, current PSC-derived HLCs exhibit features that are more consistent with fetal hepatocytes, a phenomenon also observed for other PSC-derived cell types [14]. The observation that more mature HLCs can be generated from PSCs via timed upregulation of HNF6 and PROX1 [15] or enforced adenovirus vector-mediated expression of ATF5, CEBPA and PROX1 [16], suggests that the immature phenotype is at least in part due to the suboptimal expression of key hepatic transcription factors. Mapping gene expression profiles and transcription factor motifs that define the differentiation of stem cells into HLC [17] further supports the idea that key stimuli required to induce, or maintain, appropriate gene expression in mature hepatocytes are lacking from current protocols[18]. Previously random, inefficient screens have been undertaken to identify small molecules that can promote hepatic specification, proliferation and maturation, [19, 20]. We hypothesized that comparing the transcriptomes of HLC from sequential stages of *in vitro* differentiation with RNA sequencing data from human fetal liver [21] and adult hepatocytes [22], would be a more direct approach to identify suboptimal gene regulatory networks responsible for later stages of hepatocyte differentiation and maturation. Using upstream pathway prediction analyses we here predict and subsequently experimentally validate that thyroid hormone triiodothyronine (T3) and TGF-β receptor inhibitor ALK5i II upregulate the expression of the sub-optimally expressed key transcription factors CEBPA and HNF4A, and drive the acquisition of a more mature hepatocyte phenotype, including expression of CYP3A4 and other cytochrome P450 enzymes involved in xenobiotics metabolism.

## Material and Methods

### Pluripotent stem cell-hepatocyte-like cell (PSC-HLC) differentiation

HUES7 human embryonic stem cells (ESC) were maintained on inactivated mouse embryonic feeder cells (MEFs) as previously described, using TrypLE Express (Thermo Fisher Scientific, US) as a dissociation agent for passaging [13]. Definitive endoderm differentiation commenced 3-4 days post-passage onto fresh MEFs using 25 ng/ ml WNT3a (R&D Systems, UK) and 100 ng/ ml Activin-A (Peprotech, UK), diluted in RPMI media (Sigma-Aldrich, UK) containing 0.5% FBS (Thermo Fisher Scientific) days 1-2 (Stage 1A), and without WNT3a days 5-8 (Stage 1B). Days 5-10 (hepatoblast differentiation/ Stage 2) switched to complete Hepatocyte Culture Medium (HCM) (consisting of HBM Basal Medium and HCM SingleQuot Supplements (Lonza, UK)) containing 20 ng/ ml BMP2 and 30 ng/ ml FGF2 (both Peprotech). Hepatoblast specification/ Stage 3A commenced on days 11-15 with 20ng/ ml HGF (Peprotech) diluted in complete HCM, followed by maturation/ Stage 3B during days 16-27 with 30 ng/ ml OSM (Peprotech) and 100 nM Dexamethasone (Sigma-Aldrich, UK) in complete HCM.

For the feeder-free experiments, H9 ESCs were maintained on Matrigel™ hESC-Qualified Matrix (Corning), in mTeSR™ Plus (STEMCELL Technologies) and passaged using 0.5 mM EDTA (Thermo Fisher Scientific). For HLC differentiation cells were seeded onto Matrigel™ coated 24-well tissue culture plates at 130,000 cells/ cm^2^ in mTeSR™ Plus containing 10 μM ROCK inhibitor (STEMCELL Technologies). After 24 hours the differentiation protocol was commenced using STEMdiff™ Definitive Endoderm Kit (STEMCELL Technologies) days 1-4 (Stage 1.I) following manufacturers guidelines, followed by 100 ng/ ml Activin-A (Peprotech) and 2% B-27™ Supplement minus insulin (Thermo Fisher Scientific) diluted in RPMI 1640 Medium (Thermo Fisher Scientific) days 5-8 (Stage 1.II). For hepatoblast differentiation/ Stage 2, 10 ng/ ml FGF10 and 10 ng/ ml BMP4 was diluted in complete HCM for days 9-12. Hepatocyte specification and maturation involved 50 ng/ml HGF (all Peprotech) and 30 ng/ml OSM (In Vitro Technologies) diluted in complete HCM for days 13-18 (Stage 3.I) and days 19-28 (Stage 3.II).

### RNA sequencing, mapping and quantification

RNA-sequencing and initial data processing and analysis was undertaken using methods fully described in Gerrard et al. [21]. In brief, total RNA was isolated from pluripotent ESCs or cells at differentiation stages 1B, 2, 3A and 3B of the MEF-based differentiation protocol, using an RNeasy Mini Kit (Qiagen) and subjected to DNAse (Sigma Aldrich) treatment, according to the manufacturer’s guidelines. Experiments were run in duplicate to generate 2 samples for all stages, with the exception of hepatoblast/ Stage 2 cells, which include a third, isolated experimental sample. cDNA was generated using reverse transcriptase High Capacity RNA-to-cDNA kit (Applied Biosystems, Life Technologies) according to the manufacturer’s guidelines. cDNA samples were processed and RNA-sequenced on a Illumina HiSeq® 2000 Sequencing platform (Illumina Inc, San Diego, CA, USA), according to the manufacturer’s instructions, at the Genomic Technologies Core Facility, University of Manchester, UK. RNA-sequencing (RNAseq) reads were mapped to the human reference genome (hg19) using TopHat version 2.0.9 [23]. Gene-level counts were generated for the GENCODE18 [24] annotation using Partek (version 6.6; Partek Inc., St.Louis, MO, USA). Counts were quantile normalised across samples using the R package preprocessCore [25]. The R package EdgeR [26] was used to test for differential expression between genes at successive stages of the differentiation. For comparison, RNAseq data from fresh hepatocytes (F_Hep), HepG2 cells, undifferentiated Human embryonic fibroblasts (HEF), ESC-derived HLCs (ES_Hep) and reprogrammed fibroblast derived HLCs (iHep) (Du et al [22] from GEO Acc:GSE54066) as well as from a range of in-house analysed human embryonic tissues including liver (available as described in source reference [21]), were quantile normalised, re-mapped and counted as above.

### Principal components analysis

Principal components (PC) analysis (PCA) was performed in R (R Development Core Team) using the set of genes detected in all samples. Due to the in-experiment normalization of mRNA intensities, PCA was performed on the covariance matrix without further scaling of RNA intensities, as described in Gerrard et al. [21]. The importance of each sample in generating each PC was expressed as a loading. The position of each gene mapped onto each PC was expressed as a score. A small subset of PCs contained a large proportion of the variance in the data. Sample loadings were consistent for samples of the same biological origin. We used the R package topGO [27] to test for gene set enrichment of Gene Ontology (GO) categories at either end of each of the first two principal components (PCs). In each case, a one-tailed Wilcoxon rank sum test was applied using the ‘elimination’ algorithm to reduce redundant testing within groups of nested GO terms.

### Ingenuity® Pathway Analysis

Genes that were significantly differentially expressed (false discovery rate [FDR]= ≤ 0.001) between fresh adult hepatocytes and Stage 3B HLCs were imported into QIAGEN’s Ingenuity® Pathway Analysis (IPA®, QIAGEN Redwood City,www.qiagen.com/ingenuity) software and analysed with the core analysis function. The activity of canonical pathways was predicted in Ingenuity® Pathway Analysis based on the significant differential expression of pathway-related genes in adult liver versus Stage 3B HLCs and Stage 3B HLCs versus Stage 3A hepatocyte specification cells.

#### HLC phenotypic analyses

Q-PCR was undertaken using SYBR® Green PCR Master Mix (Applied Biosystems, UK) or SsoFast EvaGreen Supermix (Bio-Rad) following manufacturers guidelines. A list of primers used is shown in Supplementary Table S1. Q-PCR was conducted on the CFX384 Touch Real-Time PCR Detection System – (Bio-Rad) maintaining individual biological repeats with control on one experimental run. Following internal normalisation using housekeeper genes, all expression values were presented as fold change versus no treatment control. The following assays were undertaken as previously described [1, 13]. Albumin secretion was quantified using the Human Albumin ELISA Quantitation Set (Bethyl Laboratories Inc) and CYP3A4 activity was determined using P450-Glo CYP3A4 Assay with Luciferin-IPA (Promega), following the manufactures guidelines. Albumin and CYP3A4 activity were internally-normalised against protein content of each individual well, as determined by a BCA protein assay kit (Thermo Fisher Scientific) or a Bradford protein assay (Bio-Rad). For Western Blot, cells were lysed with RIPA buffer containing protease and phosphatase inhibitors. Samples were resolved under reducing conditions using denaturing TGS (tris-glycine/SDS) buffer-based polyacrylamide gel electrophoresis (SDS-PAGE). Following a wet transfer (Tris-glycine methanol buffer) to nitrocellulose membranes, blots were blocked in 5% milk and probed with selected primary antibodies (Supplementary Table S1) at 4°C overnight. Appropriate HRP-conjugated secondary antibodies were detected using Clarity ECL (BioRad). Captured images were analyzed using Image Lab 4.1 (Bio-Rad, USA) software, and adjusted background values for bands of interest were used to quantify internally-normalised expression levels for each protein lysate sample. A Mann Whitney test was used to generate statistical significance of directional changes in gene expression, Albumin secretion, CYP3A4 activity and protein expression, comparing treated cells to the vehicle control.

## Results

### *HNF4A* and *CEBPA* are under-expressed during the final stages of hepatocyte differentiation

Pluripotent human embryonic stem cells (HUES7) were directed towards HLCs by applying the protocol described by Baxter et al. 2015 (Fig. 1A), which involves timed exposure to a set of widely used soluble additives [2-4] to foster the specification of definitive endoderm-like cells (Stage 1B), hepatoblast-like cells (Stage 2), early hepatocyte-like cells (Stage 3A), and hepatocyte-like cells (Stage 3B). Quantification of mRNA abundance at each of these stages with RNAseq revealed the expected loss of pluripotency genes (*NANOG, POUF1*), transient upregulation of primitive streak markers (*HHEX, EOMES, GSC*) and induction of definitive endoderm markers (*SOX17, CXCR4, FOXA2* and *NODAL*) by the end of Stage 1B (Fig 1B). Results are consistent with our prior findings that 84-96% of cells contained nuclear transcription factors FOXA2, SOX17 and GATA4 at this Stage [13]. By Stage 2, expression of genes encoding hepatic transcription factors *HNF4A* and *CEBPA* and fetal hepatic genes such as *AFP* and *DLK1* increased consistent with the formation of hepatoblast-like cells. At this Stage around two thirds of cells strongly express nuclear HNF4A and 91% of cells stain positive for AFP protein, as previously shown [13]. During the final phases of hepatocyte specification (Stage 3A) and early maturation (Stage 3B) expression of genes encoding proteins for phase I (*CYP3A7, CYP3A4, CYP2C9, ADH1A*) and phase II (*UGT2B7, UGT1A1*) xenobiotic metabolism were upregulated. While this overall pattern of gene expression was encouraging of accurate hepatocyte differentiation, as indicated by robust *ALB* gene expression and protein secretion in HLCs [13], we noticed that mRNA expression levels of both *HNF4A* and *CEBPA* decreased markedly by Stage 3B (Fig. 1B), a phenomenon also observed with other protocols [3]. This loss of *HNF4A* in HLCs at Stage 3B reflected a failure to sustain the switch in promoter activity from upstream (termed developmental, P2) to proximal (termed mature, P1) elements (Supplementary Fig. S1) [28, 29]. We hypothesised that this reduced expression of *HNF4A* and *CEBPA* would likely compromise HLC maturation and was due to under-representation of important gene networks normally active in mature adult hepatocytes. To identify such gene networks we computationally compared the transcriptomes of our *in vitro* generated HLCs with RNAseq data sets from human embryonic liver development [21] and a separate study of hepatic differentiation and human adult hepatocyte sequencing [22].

**Figure 1.**
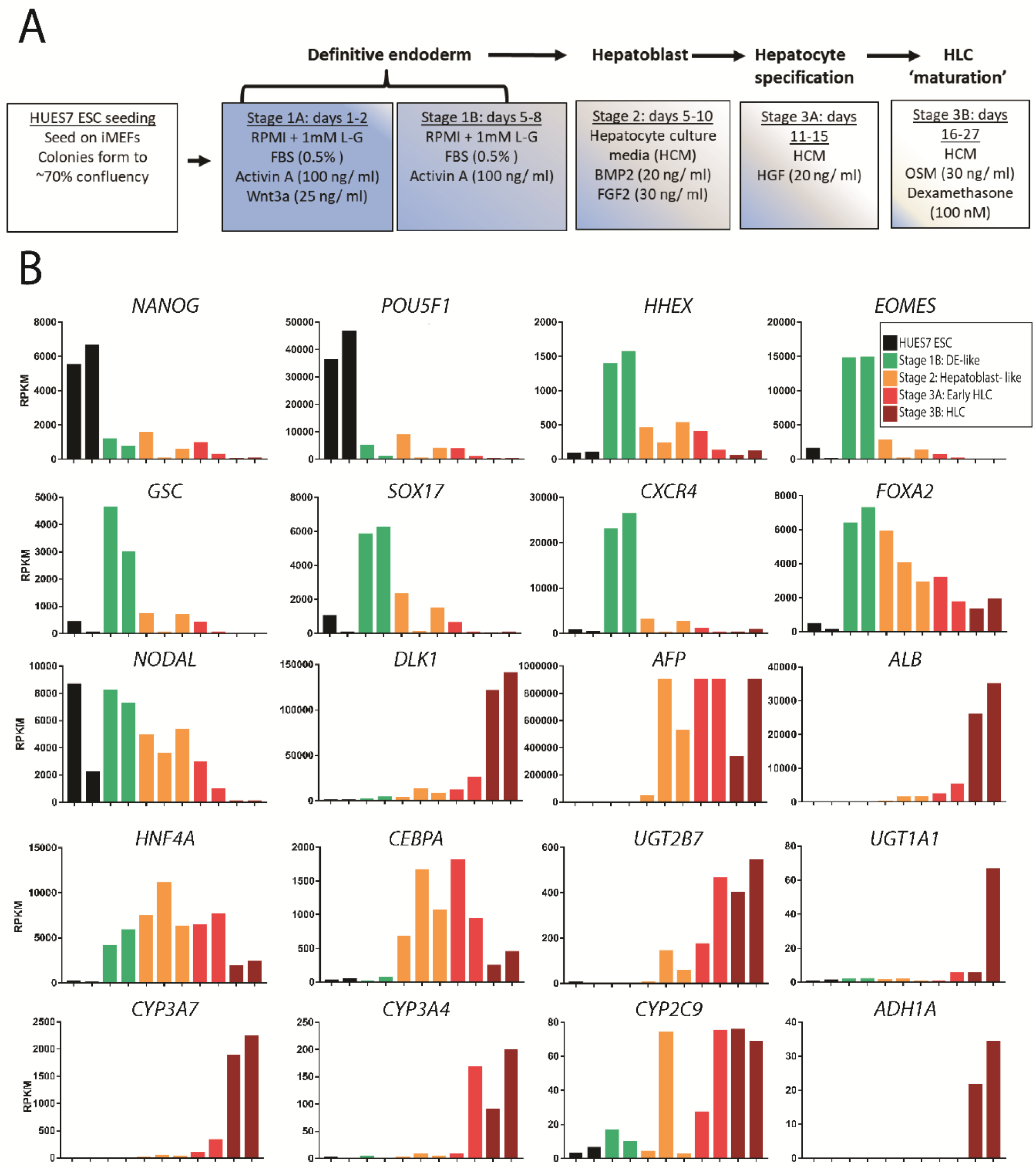
HLC differentiation mimics in vivo liver development. (A) Original hepatocyte-like cell (HLC) differentiation protocol from Baxter et al. 2015. (B) Normalised RNA sequencing expression profiles for selected genes of interest at four stages of HLC differentiation. Coloured bars represent the biological replicate samples for each stage. RPKM, reads per kilobase of transcript per million mapped transcripts; ESC, embryonic stem cell; iMEFs, inactivated mouse embryonic feeder cells; FBS, fetal bovine serum; HCM, hepatocyte culture media; DE, definitive endoderm; HLC, hepatocyte-like cell.

### Bioinformatic analysis identifies gene regulatory networks that are under-represented in HLCs compared to adult cells

We integrated our RNAseq data from cells at each stage of *in vitro* differentiation with our replicated data from seven tissues during human organogenesis, including the liver [Carnegie stages 14-20 (∼week 5-8)] [21], fresh adult human hepatocytes (F_HEP), the hepatocellular carcinoma cell line HEPG2 cells, and human embryonic fibroblasts (HEF) [22]. We undertook Principle Components Analysis (PCA) (Fig. 2A). PC1, which accounted for almost 30% of variance of the total dataset, clearly demonstrated the progression from undifferentiated PSCs (positive PC1 loadings) through differentiation stages 1 and 2 to HLCs with negative PC1 loadings. Replicates were by and large clustered closely together, albeit that the three Stage 2 hepatoblast-like populations were positioned at different points between the transitions from Stage 1B to Stage 3 populations. Two further features could be observed. Only hepatocyte-derived, HLC populations and 2 of the 3 hepatoblast-like populations had strongly negative PC1 loadings (shaded region, Fig. 2A), which by gene ontology analysis was highly enriched for numerous hepatocyte functions (Fig. 2B). In addition, PC2 separated PSC-derived HLCs (negative loadings) from fresh adult hepatocytes (strongly positive loadings) with fetal liver and HepG2 cells positioned in-between. Genes with positive PC2 loadings revealed enrichment for Gene Ontology (GO) terms indicative of mature hepatocyte function related to the metabolism of xenobiotics, fatty acids, steroids and mitochondrial function (Fig. 2B). 3767 of the 7305 differentially expressed genes (false discovery rate [FDR] = ≤ 0.001) between adult hepatocytes and HLCs were sub-optimally expressed in Stage 3B HLCs. Approximately one third (1270) of these were shared with fetal liver compared to adult hepatocytes (Supplementary Fig. S2). Proportions were similar when only considering transcription factors. Taken together, these analyses were reassuring that at a genome-scale our HLC differentiation was effective in producing hepatocyte-like differentiation, and not transition to a range of alternative cell fates. However, we had also discovered large-scale under-representation of gene expression in HLCs compared to adult hepatocytes that aligned to key aspects of phenotypic maturity.

**Figure 2.**
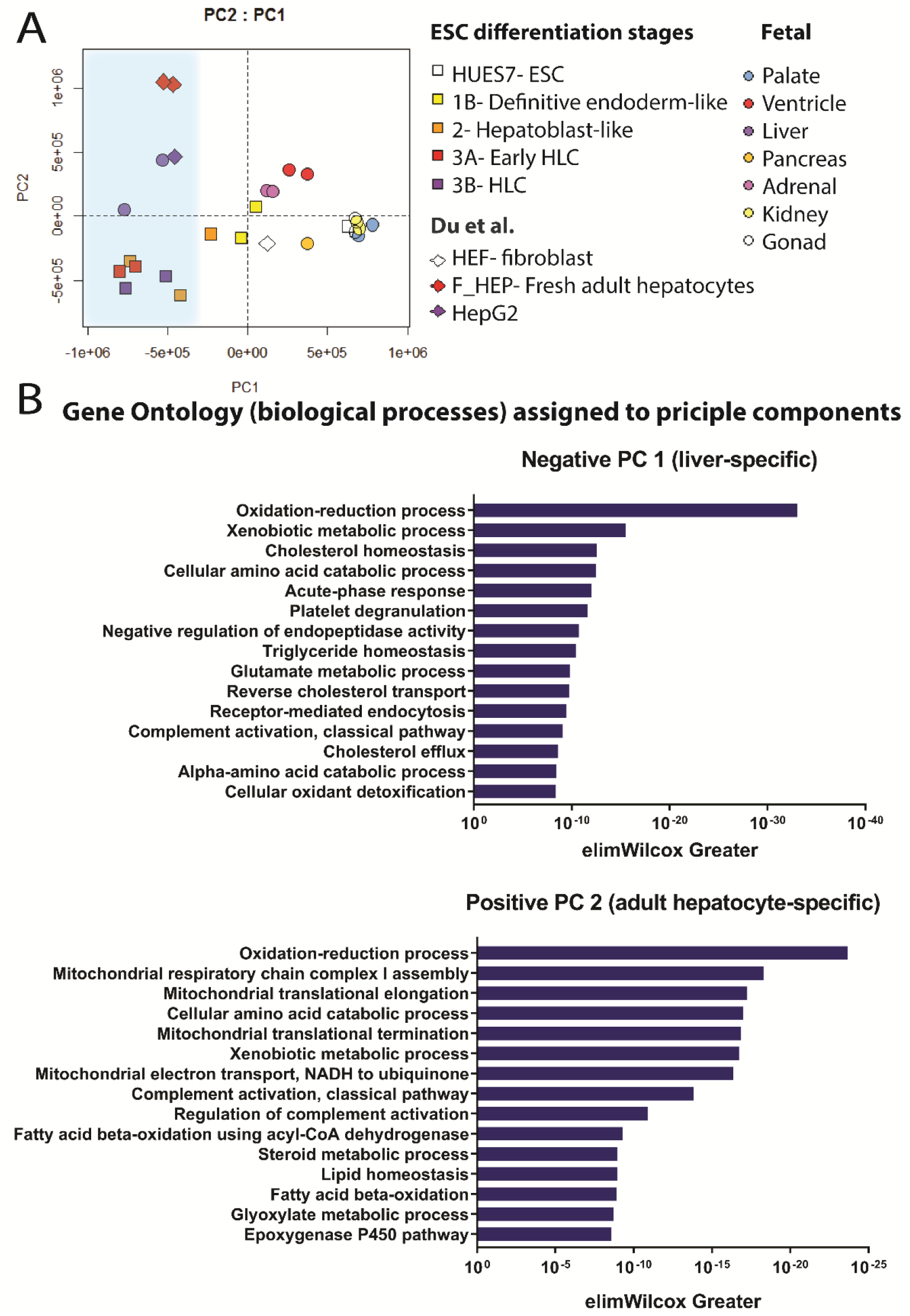
Principle components analysis identifies adult liver-specific processes. (A). Principle components analysis (PCA) of cell populations from the HLC differentiation process including additional datasets from Du et al. 2014 and Gerrard et al. 2016. The shaded area to the left hand side is to illustrate the hepatocyte and hepatocyte-like datasets typified by negative PC1 loadings. (B) Top 15 gene ontology (GO) terms associated with a negative principle component (PC) 1 loading (top panel) and a positive PC 2 loading (bottom panel).ESC, embryonic stem cell; HLC, hepatocyte-like cell.

### Pathway analysis identifies signalling that is activated but under-represented in HLCs

Rather than simply identify missing hepatocyte functions, we wanted to discover gene pathways that might be able to be manipulated to drive further maturation of HLCs towards adult hepatocytes. We reasoned three scenarios: some pathways at least partially activated in HLCs might require further upregulation; some pathways, active during HLC formation and early maturation, might require silencing; and other pathways might be down-regulated in HLCs but require upregulation in adult hepatocytes. Therefore, we applied the Canonical Pathways predictor function within Ingenuity Pathway Analysis (IPA) software on the genes differentially expressed between fresh adult hepatocytes (F_HEP) and HLCs (Stage 3B), and between HLCs (Stage 3B) versus early HLCs (Stage 3A). Two findings were immediately apparent. Firstly, some pathways, such as acute phase response signalling, were not statistically different between HLCs and adult hepatocytes, suggesting maturation of some gene regulatory pathways is possible *in vitro* with current protocols. In contrast, other liver-specific pathways, such as the complement system, coagulation system, FXR/RXR activation and xenobiotic metabolism signalling, were upregulated during the formation and early maturation of HLCs, but were deficient when compared to adult hepatocytes (Fig. 3A). Both of these scenarios could be visualised by heatmap where acute phase response pathways in HLCs had profiles similar to adult hepatocytes (Supplementary Fig. S2A) whilst the latter were more similar to embryonic liver (Supplementary Fig. S2B-D). Some pathways, such as TGF-b signalling, activated during HLC formation and early maturation, are markedly inhibited in adult hepatocytes (Fig. 3B); and finally, other pathways, such as the LXR/RXR signalling, were significantly inhibited during HLC formation and early maturation, but are active in adult hepatocytes (Fig. 3C). These combined findings led us to hypothesize that manipulating these differentially active signalling pathways (particularly those with opposing expression patterns to adult liver) might improve HLC maturation.

**Figure 3.**
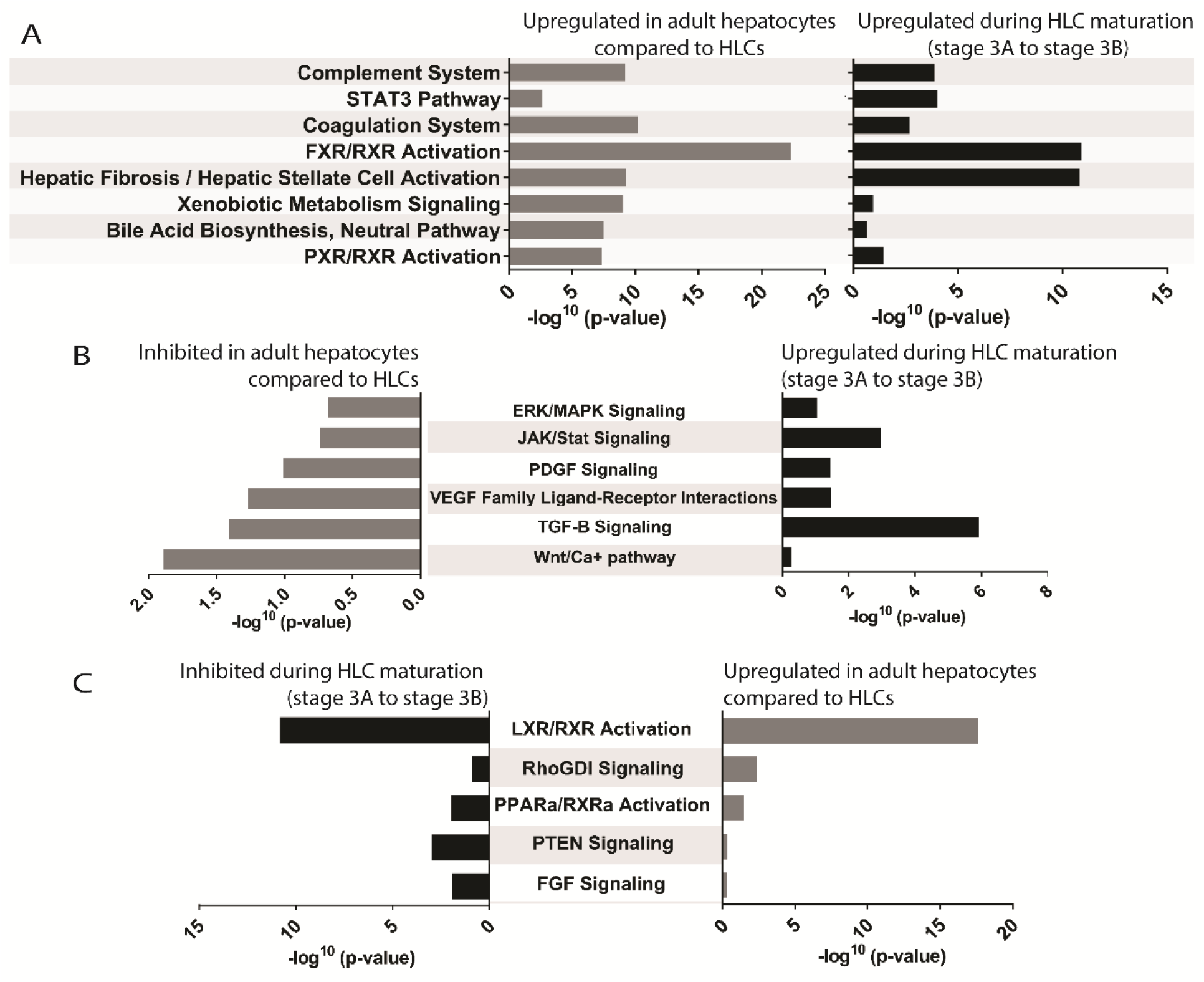
Identifying adult liver-specific canonical pathways to target. (A) Top canonical pathways upregulated in adult hepatocytes compared to hepatocyte-like cells (HLCs) as well as upregulated during the attempted maturation of HLCs (Stage 3B) versus early HLCs (stage 3A). (B) Signalling pathways upregulated during the attempted maturation of HLCs but inhibited in adult hepatocytes. (C) Signalling pathways inhibited during the attempted maturation of HLCs but active in adult hepatocytes.

To look in further detail at the pathways and how we might manipulate them, we applied the Upstream Regulator Analytic function within Ingenuity® Pathway Analysis (IPA®) software to those genes which were significantly over-expressed in adult hepatocytes versus HLCs (FDR <0.001) (Table 1A). This allowed us to impute the transcriptional regulators (TRs) PPARA, HNF1A, CEBPA and HNF4A, as ordered by p-value. HNF4A was predicted to upregulate the largest number (665) of differentially expressed genes. The computation also allowed the identification of compounds having the same effect (Table 1C). Interestingly, dexamethasone was the most significant compound predicted to upregulate the largest number of differentially expressed genes (700), a synthetic glucocorticoid already widely used in HLC maturation steps, including our protocol. Amongst the new candidates, we identified the thyroid hormone L-triiodothyronine (T3), which was predicted to upregulate 158 genes sub-optimally expressed in HLCs compared to adult hepatocytes. We were attracted to this finding for a number of reasons: thyroid hormone is known to induce LXR expression in hepatic cells [30] and so it was consistent with our earlier identification of under-represented LXR/RXR signalling in HLCs compared to adult hepatocytes (Fig. 3C); both thyroid hormone and LXR are key regulators of lipid metabolism [31]; and GO terms for lipid metabolism (lipid homeostasis, fatty acid beta-oxidation and the glyoxylate metabolism) were all identified as missing in HLCs and are positive discriminators of an adult hepatocyte phenotype from the PCA (Fig. 2B). In agreement with this, GO and KEGG pathway analyses of the 158 differentially expressed genes predicted to be targeted by T3 (Table 1C) identified many terms for lipid and cholesterol metabolic processes (Table 2A and B). The top KEGG pathway was PPAR signalling, in keeping with the observation that PPARA and PPARG are amongst the top transcriptional regulators of adult hepatocytes versus HLCs (Table 1A; 227 and 170 genes respectively). In rat, T3 increases hepatic glutamine synthetase (GS) activity [32], a marker of pericentral hepatocytes particularly important for xenobiotic metabolism [33, 34]. For this combination of reasons, we carried T3 forward as a candidate for maturing HLC.

**Table 1.** Upstream regulators of adult liver-specific genes.

| Upstream Regulator | Fold Change | p-value of overlap | Activation z-score | No. DEGs Targeted | Molecule Type |
| --- | --- | --- | --- | --- | --- |
| PPARA | 2.651 | 1.29E-40 | 2.626 | 227 | ligand-dependent nuclear receptor |
| HNF1A | 1.569 | 1.38E-26 | 6.119 | 210 | transcription regulator |
| CEBPA | 6.209 | 3.55E-23 | 2.655 | 194 | transcription regulator |
| HNF4A | 3.014 | 7.59E-19 | 7.967 | 665 | transcription regulator |
| ACOX1 | 2.999 | 4.01E-17 | 2.168 | 83 | enzyme |
| PPARG | 0.114 | 2.52E-14 | 4.724 | 170 | ligand-dependent nuclear receptor |
| PPARGC1A | 2.670 | 1.17E-13 | 6.113 | 100 | transcription regulator |
| AHR | 1.061 | 1.66E-13 | 3.566 | 140 | ligand-dependent nuclear receptor |
| INSR | 1.226 | 2.28E-10 | 3.317 | 141 | kinase |
| NR1I3 | 9.605 | 3.32E-08 | 3.145 | 51 | ligand-dependent nuclear receptor |

**B** Over-expressed in HLCs and (predicted) inhibited in adult hepatocytes versus HLCs
| Upstream Regulator | Fold Change | p-value of overlap | Activation z-score | No. DEGs Targeted | Molecule Type |
| --- | --- | --- | --- | --- | --- |
| TGFB1 | -4.099 | 1.28E-48 | -7.571 | 658 | growth factor |
| ERBB2 | -0.101 | 1.12E-34 | -5.413 | 299 | kinase |
| SP1 | -0.644 | 3.40E-24 | -3.313 | 231 | transcription regulator |
| CTNNB1 | -0.309 | 1.05E-17 | -4.665 | 235 | transcription regulator |
| MYC | -2.901 | 5.39E-14 | -3.388 | 349 | transcription regulator |
| HGF | -1.379 | 6.31E-13 | -5.514 | 192 | growth factor |
| EGF | -9.588 | 1.14E-12 | -5.585 | 188 | growth factor |
| TWIST1 | -6.367 | 3.94E-12 | -2.763 | 75 | transcription regulator |
| FGF2 | -0.358 | 7.54E-12 | -3.733 | 141 | growth factor |
| STAT3 | -0.507 | 1.65E-10 | -2.202 | 175 | transcription regulator |

**C** Exogenous and endogenous chemicals (predicted) active in adult hepatocytes versus HLCs
| Upstream Regulator | p-value of overlap | Activation z-score | No. DEGs Targeted | Molecule Type |
| --- | --- | --- | --- | --- |
| dexamethasone | 5.54E-42 | 2.617 | 700 | chemical drug |
| pirinixic acid | 5.51E-26 | 4.906 | 218 | chemical toxicant |
| TO-901317 | 1.13E-20 | 2.27 | 125 | chemical reagent |
| PD98059 | 2E-14 | 4.546 | 225 | chemical - kinase inhibitor |
| rosiglitazone | 1.25E-12 | 5.74 | 176 | chemical drug |
| L-triiodothyronine | 2.07E-12 | 3.27 | 158 | chemical - endogenous mammalian |
| fenofibrate | 3.43E-12 | 3.063 | 119 | chemical drug |
| methotrexate | 4.35E-12 | 2.357 | 113 | chemical drug |
| clofibrate | 1.38E-11 | 3.086 | 57 | chemical drug |
| GW 4064 | 1.63E-11 | 2.209 | 38 | chemical toxicant |

**Table 2.**
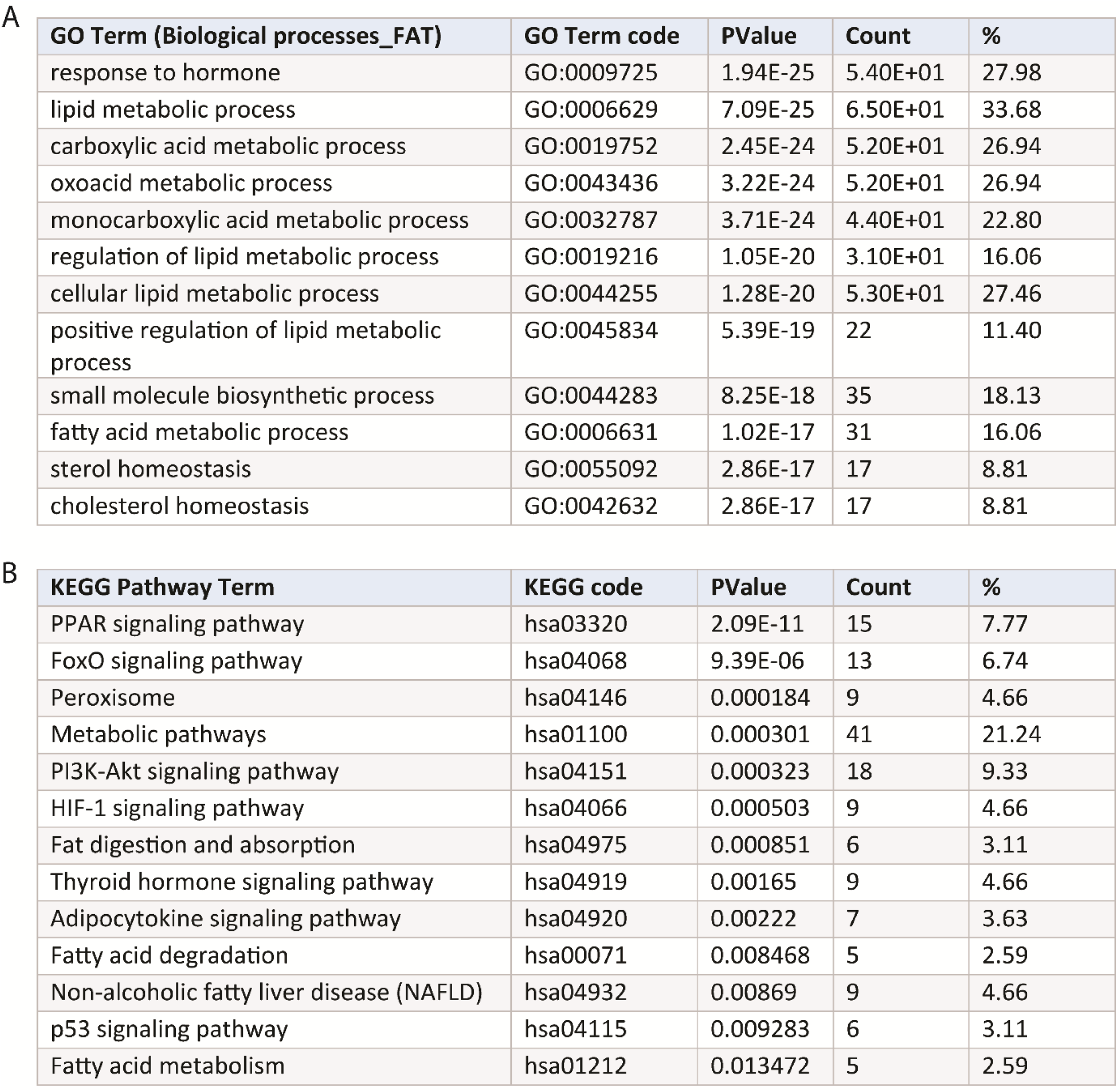
GO terms and KEGG pathway analysis of adult liver-specific genes regulated by T3.

| GO Term (Biological processes_FAT) | GO Term code | PValue | Count | % |
| --- | --- | --- | --- | --- |
| response to hormone | GO:0009725 | 1.94E-25 | 5.40E+01 | 27.98 |
| lipid metabolic process | GO:0006629 | 7.09E-25 | 6.50E+01 | 33.68 |
| carboxylic acid metabolic process | GO:0019752 | 2.45E-24 | 5.20E+01 | 26.94 |
| oxoacid metabolic process | GO:0043436 | 3.22E-24 | 5.20E+01 | 26.94 |
| monocarboxylic acid metabolic process | GO:0032787 | 3.71E-24 | 4.40E+01 | 22.80 |
| regulation of lipid metabolic process | GO:0019216 | 1.05E-20 | 3.10E+01 | 16.06 |
| cellular lipid metabolic process | GO:0044255 | 1.28E-20 | 5.30E+01 | 27.46 |
| positive regulation of lipid metabolic process | GO:0045834 | 5.39E-19 | 22 | 11.40 |
| small molecule biosynthetic process | GO:0044283 | 8.25E-18 | 35 | 18.13 |
| fatty acid metabolic process | GO:0006631 | 1.02E-17 | 31 | 16.06 |
| sterol homeostasis | GO:0055092 | 2.86E-17 | 17 | 8.81 |
| cholesterol homeostasis | GO:0042632 | 2.86E-17 | 17 | 8.81 |

| KEGG Pathway Term | KEGG code | PValue | Count | % |
| --- | --- | --- | --- | --- |
| PPAR signaling pathway | hsa03320 | 2.09E-11 | 15 | 7.77 |
| FoxO signaling pathway | hsa04068 | 9.39E-06 | 13 | 6.74 |
| Peroxisome | hsa04146 | 0.000184 | 9 | 4.66 |
| Metabolic pathways | hsa01100 | 0.000301 | 41 | 21.24 |
| PI3K-Akt signaling pathway | hsa04151 | 0.000323 | 18 | 9.33 |
| HIF-1 signaling pathway | hsa04066 | 0.000503 | 9 | 4.66 |
| Fat digestion and absorption | hsa04975 | 0.000851 | 6 | 3.11 |
| Thyroid hormone signaling pathway | hsa04919 | 0.00165 | 9 | 4.66 |
| Adipocytokine signaling pathway | hsa04920 | 0.00222 | 7 | 3.63 |
| Fatty acid degradation | hsa00071 | 0.008468 | 5 | 2.59 |
| Non-alcoholic fatty liver disease (NAFLD) | hsa04932 | 0.00869 | 9 | 4.66 |
| p53 signaling pathway | hsa04115 | 0.009283 | 6 | 3.11 |
| Fatty acid metabolism | hsa01212 | 0.013472 | 5 | 2.59 |

The most significant imputed regulator for down-regulated genes in adult hepatocytes compared to HLCs re-identified TGFB signalling (Table 1B and Fig. 3B). TGFB was predicted to target nearly twice as many differentially expressed genes (658) compared to the other identified factors (Table 1B). Since there are different isoforms of TGFB we examined their expression during HLC differentiation and in comparison to adult hepatocytes. *TGFB3* was relatively unchanged between HLCs, fetal liver and adult hepatocytes (Fig. 4A). In contrast, *TGFB1* and *TGFB2* levels were considerably higher in HLCs compared to adult hepatocytes (14.7 and 35.5-fold, respectively). Similar fold changes (8.5 and 53-fold) were observed using data of PSC-derived HLCs from Du et al [22] (Fig. 4A). Interestingly, TGFB1 has been reported to down-regulate *HNF4A* expression in mice [35], consistent with our IPA analysis (Fig. 4B), and can also decrease HNF4A DNA binding in HEPG2 cells [36]. Therefore, alongside T3, we considered inhibition of TGFB signalling, most likely mediated through *TGFB1* or *TGFB2*, as a good candidate to improve HLC maturation.

**Figure 4.**
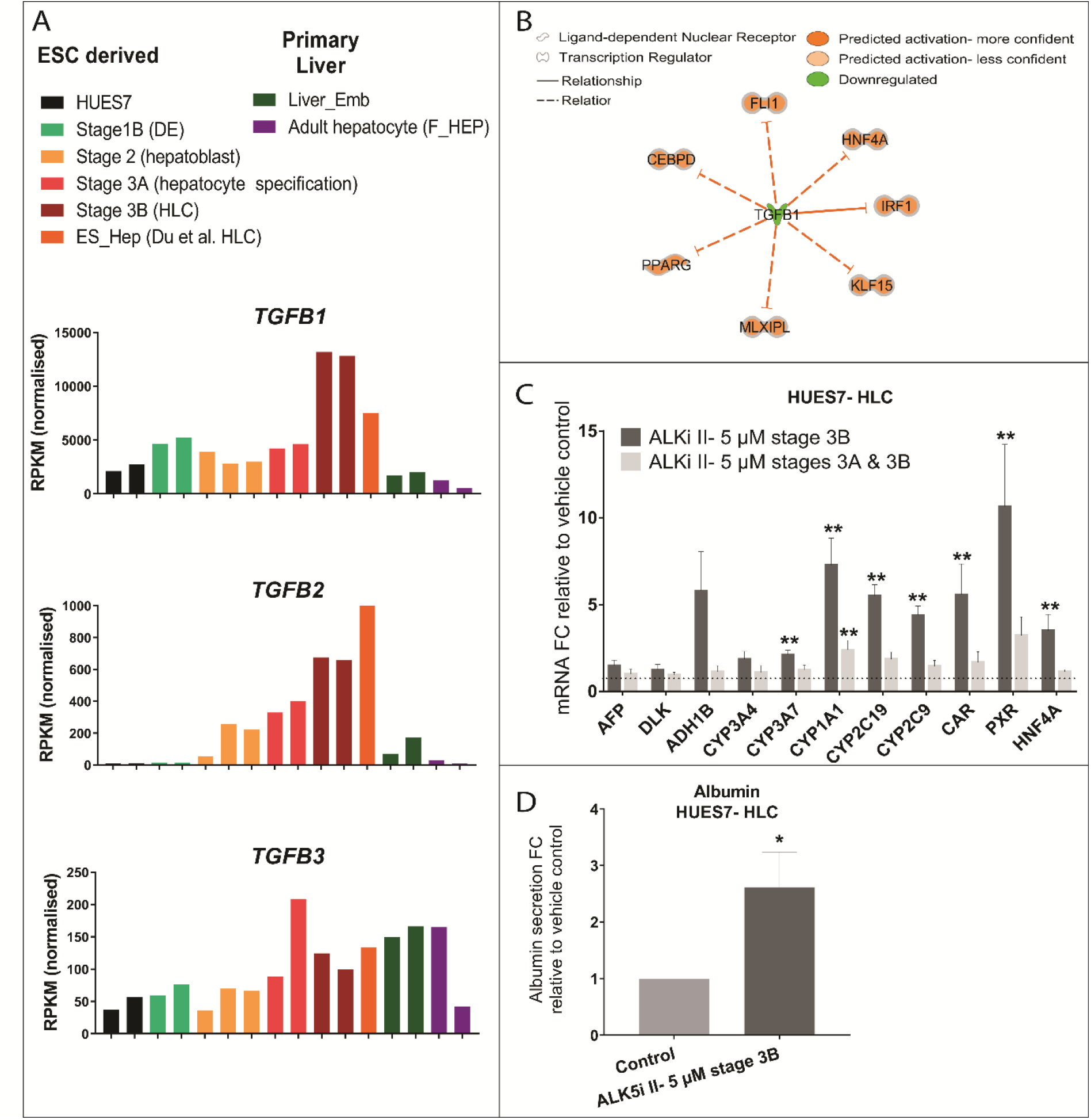
TGFB inhibition during HLC differentiation improves hepatic maturation. (A) Bar charts showing TGFB1, TGFB2 and TGFB3 transcript values (RNA sequencing) from hepatocyte like cell (HLC) differentiation stages from this investigation and from published HLC differentiation studies (ES_Hep), adult liver (both Du et al. 2014) and fetal liver/ Liver_Emb (Gerrard et al. 2016). (B) Schematic of the Ingenuity Pathway Analysis predicted relationships of TGFB1 with the adult liver-specific transcription factors sub-optimally expressed in Stage 3B HLCs compared to adult liver. (C) Bar charts showing fold increase in mRNA levels of liver-specific genes of interest (quantitative PCR, *n* = 4-5) following TGFB inhibition through the use of 5 µM ALK5i II during hepatocyte specification Stage 3A onwards or hepatocyte maturation Stage 3B alone. Fold change versus vehicle control (indicated by dotted line). (D) Bar chart showing albumin secretion fold change in HLCs treated with 5 µM ALK5i II during Stage 3B compared to the vehicle control (ELISA, *n* = 6). Albumin values normalised to protein quantity of sample. Data are presented as mean + standard error of the mean (SEM); *p < 0.05, **p < 0.01, using a Mann Whitney test.

### ALK5i II and T3 treatments improve HLC maturation

To inhibit TGFB signalling, we selected ALK5i II, a molecule that selectively targets TGFβRI kinase and inhibits signalling by all three TGFB isoforms. We added 5 µM ALK5i II either from hepatocyte specification Stage 3A onwards or only during hepatocyte maturation Stage 3B. As expected based on TGFB overexpression patterns (Fig. 4A), the latter was consistently the most effective, resulting in a 3.5-fold increase in *HNF4A* mRNA levels (3.5-fold) and significant increases in the expression of many other genes involved in xenobiotic metabolism such as: *CYP3A7* (2.1-fold), *CYP1A1* (7.3-fold increased), *CYP2C19* (5.5-fold), *CYP2C9* (4.4-fold), *Constitutive Androstane Receptor* (*CAR*) (5.6-fold), and *Pregnane X Receptor* (*PXR*) (10.6-fold) (Fig. 4C). CAR and PXR both regulate CYP450 expression. *PXR* was identified earlier by heatmap as deficient in HLCs (Supplementary Fig. S3D). ALK5i II treatment did not alter expression of fetal hepatocyte markers, *AFP* and *DLK1* but did increase albumin secretion 2.6 fold (Fig. 4D).

We next tested T3 (1 µM) in a similar manner from Stage 3A or during Stage 3B only. Broad improvements were observed. The timing of addition, either from Stage 3A or during Stage 3B only, was somewhat unimportant for a number of genes: *HLF* 8.3- and 5.7-fold, *KLF9* 3.7- and 2.9-fold, *CEBPA* 1.7- and 1.5-fold, *CYP1A1* 3.6- and 2.9-fold, and *ADH1B* 2.7- and 5.2-fold respectively for all (Fig. 5B). However, T3 treatment only for Stage 3B was required to increase expression of *CYP3A4* (3.1-fold), *CYP3A7* (2-fold), *CYP2C19* (2.1-fold) and *CYP2C9* (2.4-fold) and prevented the decrease in *CAR* expression. Neither treatment made a significant difference to *HNF4A* expression, as our IPA analysis predicted based on the relationship of T3 to the transcriptional regulators sub-optimally expressed in HLCs compared to adult liver (Fig. 5A). Albumin secretion during Stage 3B was increased 2.6-fold by T3 (Fig. 5C). While ALK5i II made little difference, 1 µM T3 also seemed to increase Glutamine synthetase (GS) protein levels in HLCs (Supplementary Fig. S4). Collectively, these data indicate that inhibiting TGFB signalling and adding T3 during the final phase of HLC differentiation enhanced a range of parameters consistent with HLC maturity.

**Figure 5.**
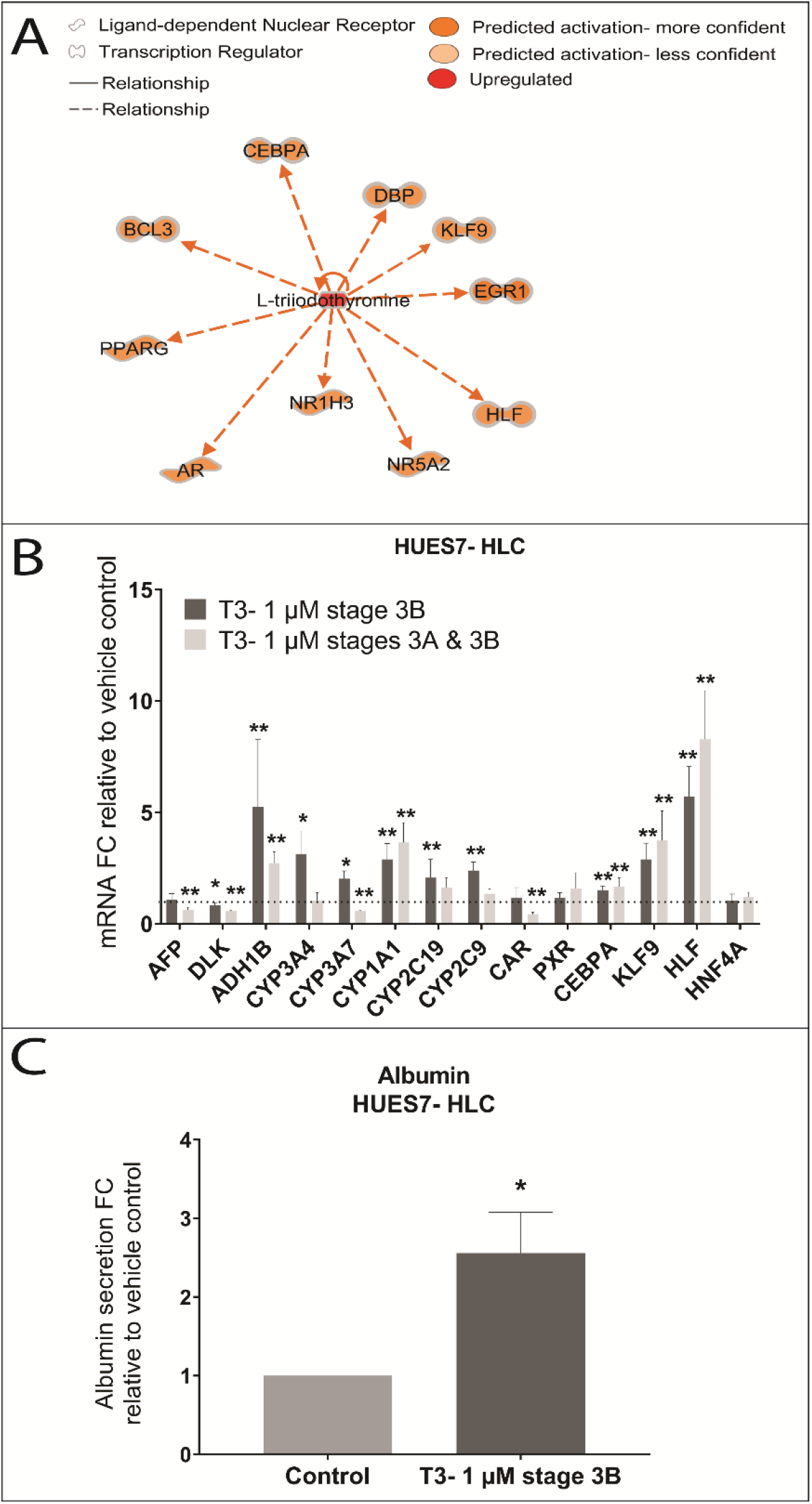
T3 treatment of HLCs improves hepatic maturation. (A) Schematic of the Ingenuity Pathway Analysis predicted relationships of Triiodothyronine (T3) with the transcription factors sub-optimally expressed in stage 3B hepatocyte-like cells (HLCs) versus adult liver. (B) Bar chart showing fold increase in mRNA levels of liver-specific genes of interest (quantitative PCR, *n* = 4– 6) following 1 µM T3 treatment during hepatocyte specification Stage 3A onward and hepatocyte maturation Stage 3B alone. Fold change compared to vehicle control (indicated by dotted line). (C) Bar chart showing albumin secretion fold change in HLCs treated with 1 µM T3 during Stage 3B compared to the vehicle control (ELISA, *n* = 4). Albumin values normalised to protein quantity of sample. Data are presented as mean + standard error of the mean (SEM); *p < 0.05, **p < 0.01, using a Mann Whitney test.

### ALK5i II and T3 also improve maturation in feeder-free hepatocyte differentiation protocols

Encouraged by these data we wanted to test our new additives in a second, potentially improved protocol. To minimize cost and enable future clinical translation, HLC differentiation protocols will ideally need to exclude animal derived products, such as mouse embryonic feeder cells (MEFS) and fetal bovine serum (FBS). Therefore, we developed a feeder-free hepatocyte differentiation method with a different ESC line (H9) based largely on Hannan et al., 2013 [11] (Fig. 6A). Cells were treated with 1 µM T3 and 5 µM ALK5i II, alone or in combination, for different lengths of time (Stage 3.II or both Stages 3.I & 3.II, identical media components). Broadly, very similar results were observed compared to addition to our monolayer protocol albeit with larger fold change improvements (Fig. 6B-C). Treatment with 1 µM T3 during Stage 3.II significantly increased transcript levels for *CEBPA* (92.1-fold), *KLF9* (517.5-fold), *HLF* (95.4-fold) and *HNF4A* (47.6-fold) and for mature liver markers *A1AT* (88.2-fold), *ALB* (24.4-fold), *CYP3A4* (114.5-fold) (Fig. 6B). T3 treatment also increased CYP3A4 activity (13.1-fold and 5.3-fold respectively for Stages 3.I and 3.I-3.II; Fig. 6F), HNF4A protein levels (2.9-fold and 2.1-fold; Fig. 6G), and HLF protein (1.4-fold and 2.4-fold respectively; Fig. 6H). Treatment with 5 µM ALK5i II for the last Stage significantly increased transcript levels of *CEBPA* (11.6-fold), *KLF9* (53.9-fold), *HLF* (104.8-fold), *HNF4A* (9.7-fold), *A1AT* (14.3-fold), *ALB* (14.1-fold), *ADH1B* (67.6-fold), *CYP2C19* (28.5-fold), *CYP2C9* (15.3-fold) and *CAR* (7.1-fold) (Fig. 6C). ALK5i II treatment also increased CYP3A4 activity (Fig. 6F), and HNF4A and CEBPA protein levels, but not HLF (Fig. 6G-I).

**Figure 6.**
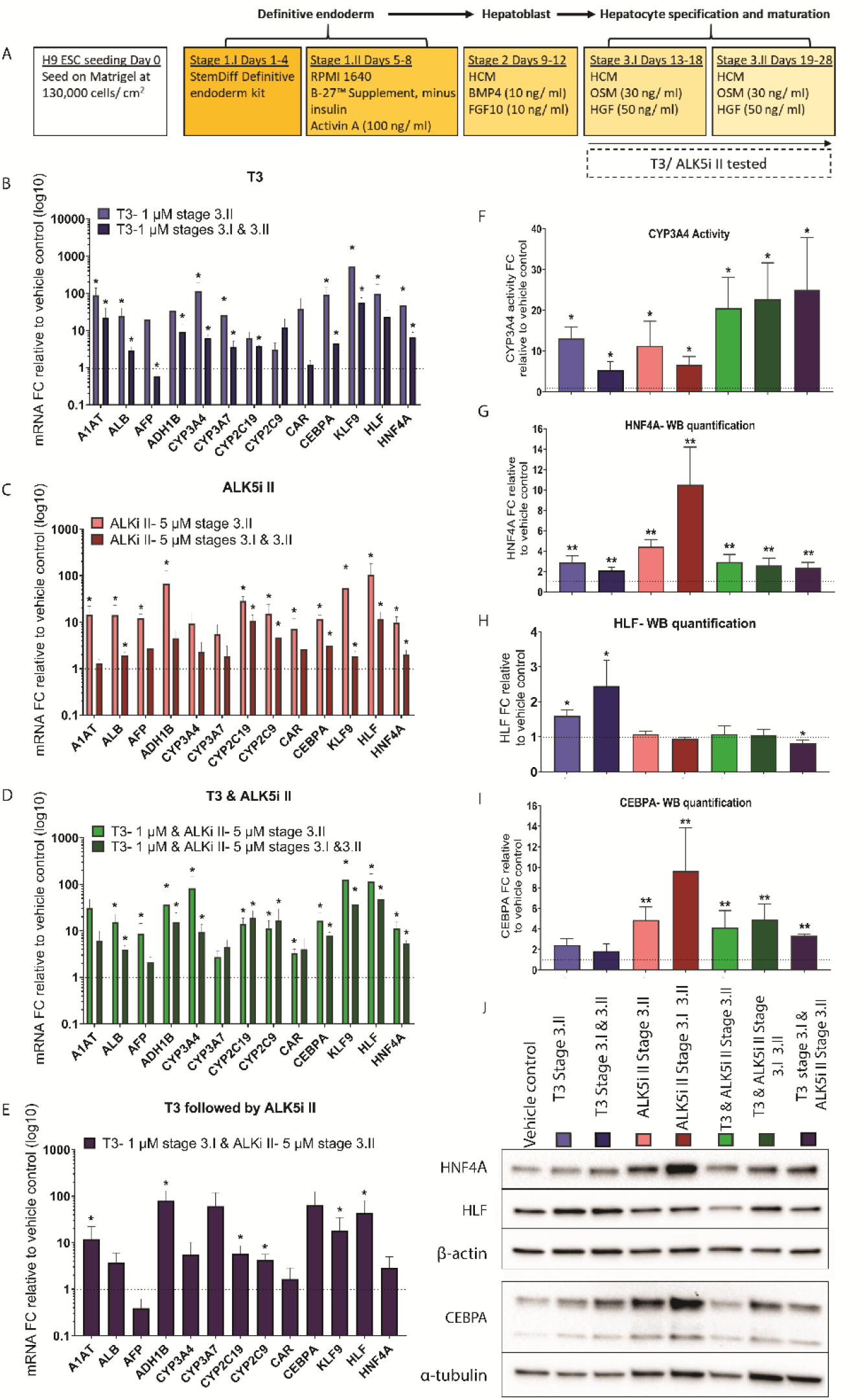
ALK5i II and T3 treatment improves feeder-free hepatocyte differentiation. (A) Feeder-free hepatocyte-like cell (HLC) differentiation protocol. (B-E) Bar charts showing fold increase in mRNA levels of liver-specific genes of interest (quantitative PCR, *n* = 4) following HLC differentiation treatment with (B) T3, (C) ALK5i II, (D) T3 and ALK5i II and (E) sequential addition of T3 followed by ALKi II during timed intervals of HLC specification and maturation (Stage 3). Duration of stage 3 treatment (3.II or 3.I and 3.II) highlighted by light and dark bars respectively. Fold change compared to vehicle control (indicated by dotted line). (F) Bar chart showing CYP3A4 activity (luciferase-IPA, *n* = 4) fold change in HLCs treated with 1 µM T3, 5 µM ALK5i II, 1 µM T3 and 5 µM ALK5i II and sequential addition of 1 µM T3 followed by 5 µM ALKi II during timed intervals of HLC specification and maturation (Stage 3), relative to vehicle control (dotted line). (G-I) Bar charts showing fold increase in protein concentration of liver-specific transcription factors (G) HNF4A, (H) HLF and (I) CEBPA following HLC differentiation treatment with 1 µM T3, 5 µM ALK5i II, 1 µM T3 and 5 µM ALK5i II and sequential addition of 1 µM T3 followed by 5 µM ALKi II during timed intervals of stage 3 (*n* = 3-6). Fold change compared to vehicle control (indicated by dotted line). Relative protein concentration determined by (J) Western Blot image quantification, normalised to B-actin or a-tubulin. Data are presented as mean + standard error of the mean (SEM); *p < 0.05, **p < 0.01, using a Mann Whitney test.

Next, we tested the two factors in combination. Increased marker gene expression was retained at broadly similar levels to treatments in isolation (Fig. 6D), as was CYP3A4 activity, and HNF4A and CEBPA protein levels (Fig. 6G-J). Sequential treatment of T3 followed by ALK5i II had little or no additional benefit on gene expression (Fig. 6E), CYP3A4 activity (Fig. 6F) or transcription factor protein levels (Fig. 6G-J).

In summary, both inhibition of TGFB signalling and thyroid hormone improved HLC maturation in two different protocols when added to the last phase of differentiation. Improvements appeared more marked in the feeder-free protocol.

## Discussion

*In vivo*, human liver maturation occurs over a period of several years and is thought to involve a complex and dynamic interplay between multiple cell types and temporally changing metabolic and xenobiotic challenges. It is therefore perhaps unsurprising that maturation of hepatocytes in stem cell derived protocols has been challenging. In this study, we compared the transcriptomes of cells during differentiation with those from primary human fetal liver and adult liver with the aim of identifying gene regulatory networks that might impact on the acquisition and/or maintenance of liver cell maturity. Our data revealed that gene regulatory pathways related to metabolism of xenobiotics, fatty acids, steroids and mitochondrial function were deficient in HLCs. We also observed that expression of *HNF4A* and *CEBPA* declined during the final phases of *in vitro* HLC generation of hPSCs. Both transcription factors are important for adult hepatocyte function and a prerequisite for liver development and adult liver regeneration [37, 38]. Computationally, we identified the inhibition of TGFB signalling and T3 as factors to enhance HLC maturity *in vitro*. Our findings that T3 can be used to promote hepatic maturation are consistent with other recent reports [39, 40]. Thyroid hormone was previously shown to both directly [41] and indirectly [42] upregulate CEBPA signalling and increases hepatic CEBPA and CEBPB levels in hypothyroid rats [43]. Thyroid hormone is also known to be important for hepatic fatty acid and cholesterol synthesis and metabolism [44]. T3 was shown to induce KLF9 in HepG2 liver cells, mouse liver, and mouse and human primary hepatocytes [45], [46] and upregulates HLF [47] in HEPG2 cells. T3 was also found beneficial in protocols to differentiate more mature pancreatic beta-cells [48]. Our data in HLCs are broadly compatible with these findings with T3 treatment of HLCs resulting in a significant increase in a broad range of hepatocyte maturity markers. The LXR/RXR pathway has been shown to regulate thyroid hormone activation [49], and both LXR and thyroid hormone receptors show similarities in molecular mechanism, target genes, and physiological roles [30, 31]. This potentially provides an explanation for why GO terms relating to the LXR/RXR pathway, lipid homeostasis, fatty acid beta-oxidation and the glyoxylate metabolic process were all sub-optimal in non-T3 treated HLCs and discriminators of an adult hepatocyte phenotype.

In contrast, we discovered that TGFB signalling was overactive in HLCs compared to adult hepatocytes, in keeping with a detrimental effect on HLC maturity. TGFB1 was previously shown to inhibit *Hnf4a* mRNA expression in mouse NMuMG cells [35] and to interfere with HNF4A function in HEPG2 cells [36]. Consistent with our bioinformatic predictions, treatment of HLCs with ALK5i II resulted in significant increases in transcript levels for mature hepatocyte-specific genes encoding CYP450 enzymes, nuclear receptors CAR and PXR. Increased albumin secretion and CYP3A4 activity were all observed. ALK5i II had no effect on the already low protein level of the biliary marker SOX9 in our HLCs. Therefore, it seems unlikely that its main effect was on altered hepatoblast fate choices known to be influenced by TGFB signalling *in vivo* [37, 50]. Instead, our observation of improvement during the last phase of two protocols strongly implies positive influence on hepatocyte maturation rather than differentiation.

In conclusion, in this study we have delineated phenotypic deficiencies in HLCs compared to mature adult hepatocytes, consistent with the relative lack of key transcription factors, HNF4A and CEBPA. We deduced and demonstrated that the addition of T3 and ALK5i II could, at least in part, reverse important aspects of these phenotypic deficiencies and result in more mature HLCs *in vitro*. Addition during the final stage of the two differentiation protocols was sufficient to produce the beneficial effects in feeder-dependent and defined feeder-free differentiation protocols and across different human pluripotent stem cell lines. Taken together, these data are encouraging and point to the substantial scope for further small molecules, metabolites and / or growth factors capable of fostering improvements in HLC function and maturity.

## Data availability statement

The authors declare that all data supporting the findings of this study are available within the article and its supplementary information files or from the corresponding author upon reasonable request. RNA-seq datasets not referenced elsewhere have been deposited in the European Genome Phenome repository [www.ebi.ac.uk/ega/home] under accession codes:

---

**Supplementary Table S1.**
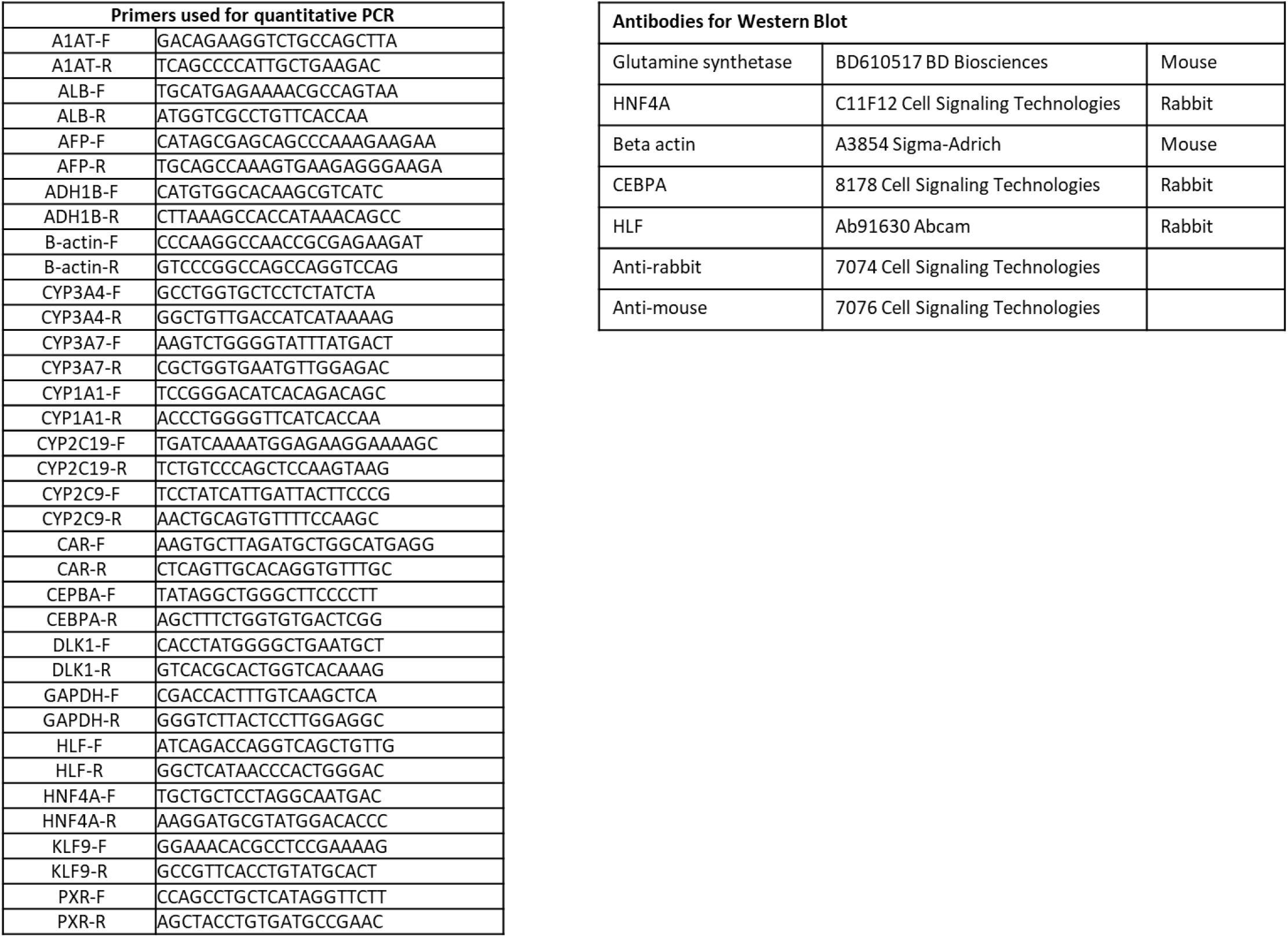
List of primers and antibodies.

**Supplementary figure S1.**
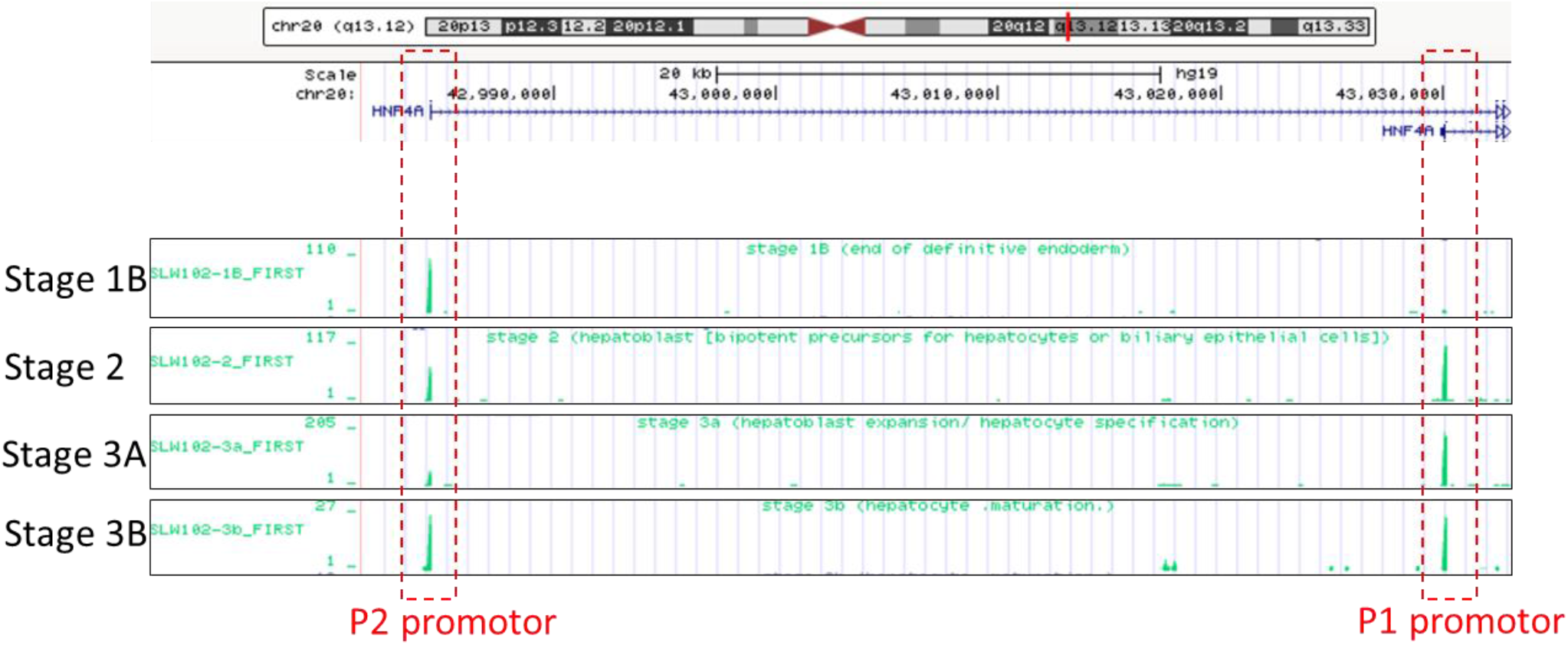
HNF4A promotor usage. HNF4A annotated using the UCSC Human Genome Browser for Stage 1B (definitive endoderm-like), Stage 2 (hepatoblast-like), stage 3A (early hepatocyte-like cell (HLC)) and Stage 3B (HLC). Red boxes highlight the shift in usage of the P2 (developmental) promotor to the P1 (mature) promoter for HNF4A, between differentiation Stages.

**Supplementary Figure S2.**
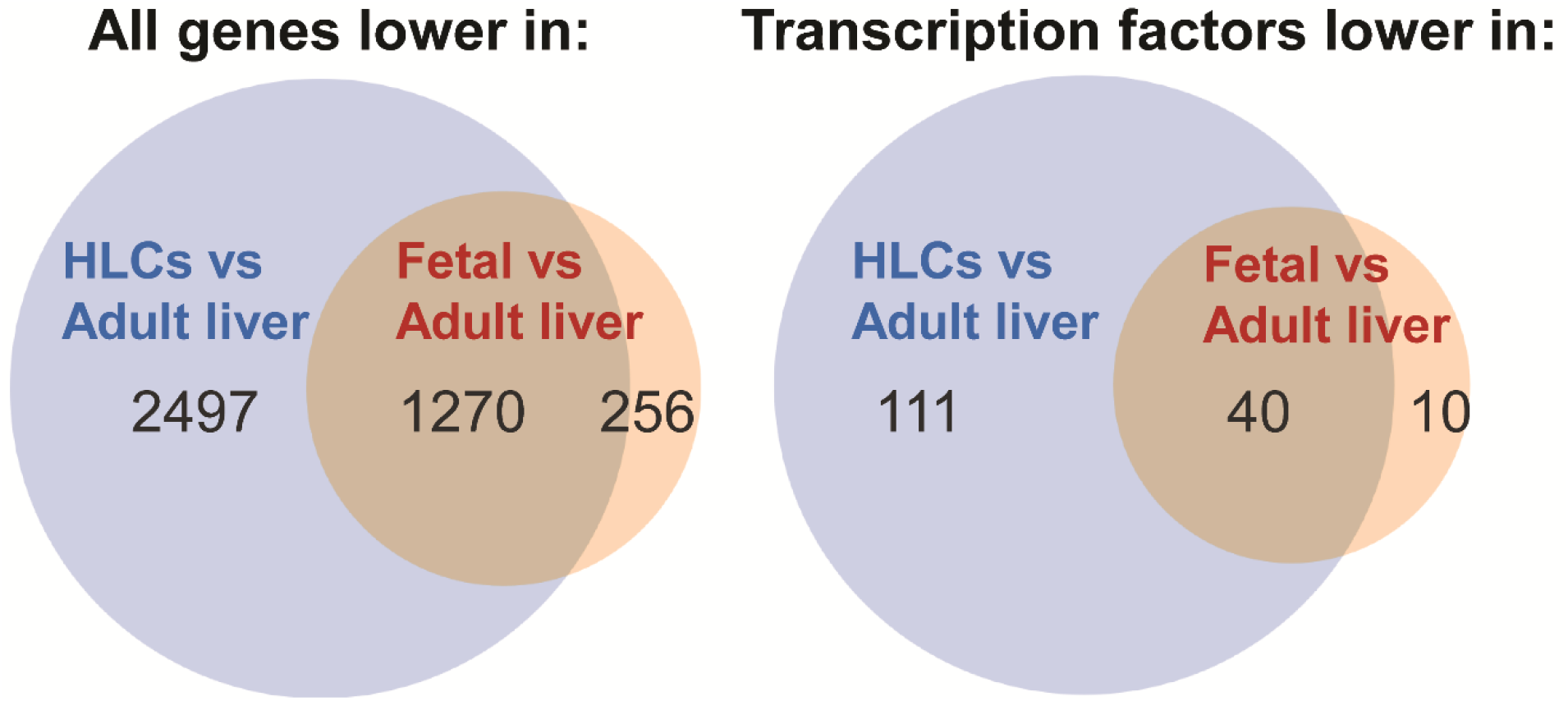
Adult liver-specific differentially expressed genes. Venn diagrams of adult liver-specific differentially expressed genes (left: all genes and right: transcription factors only) between stage 3B hepatocyte-like cells (HLCs) versus fresh adult liver from Du et al. 2014 dataset (blue) and stage 3B HLCs versus fetal liver from Gerrard et al. 2016 dataset (orange). HLC, hepatocyte-like cell

**Supplementary Figure S3.**
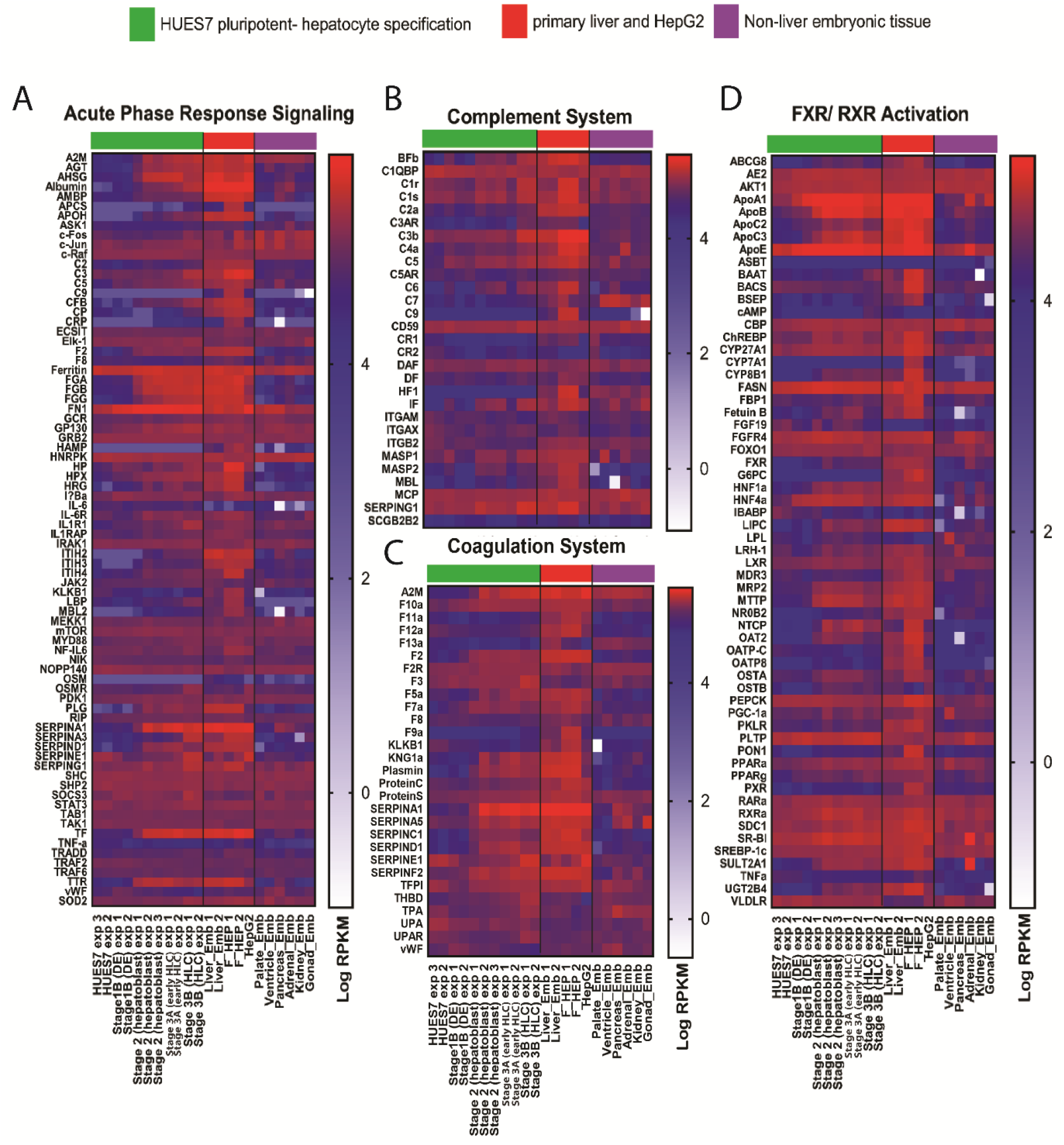
Heatmap representation of adult liver-specific canonical pathways. Normalised RNA sequencing expression profiles (log 10 RPKM values) for all Ingenuity Pathway Analysis listed genes involved in the canonical pathways (A) acute phase response signalling, (B) complement system, (C) coagulation system and (D) FXR/RXR activation, from HUES7 embryonic stem cell (ESC) line hepatocyte-like cell (HLC) differentiation stages (green bar), primary liver samples including fetal liver (liver_Emb) from Gerrard et al. 2016 and fresh adult hepatocytes (F_HEP) from Du et al. 2014 and the transformed cell line HepG2, also Du et al. 2014, (red bar), and various non-liver fetal tissue from Gerrard et al. 2016, (purple bar).

**Supplementary Figure S4.**
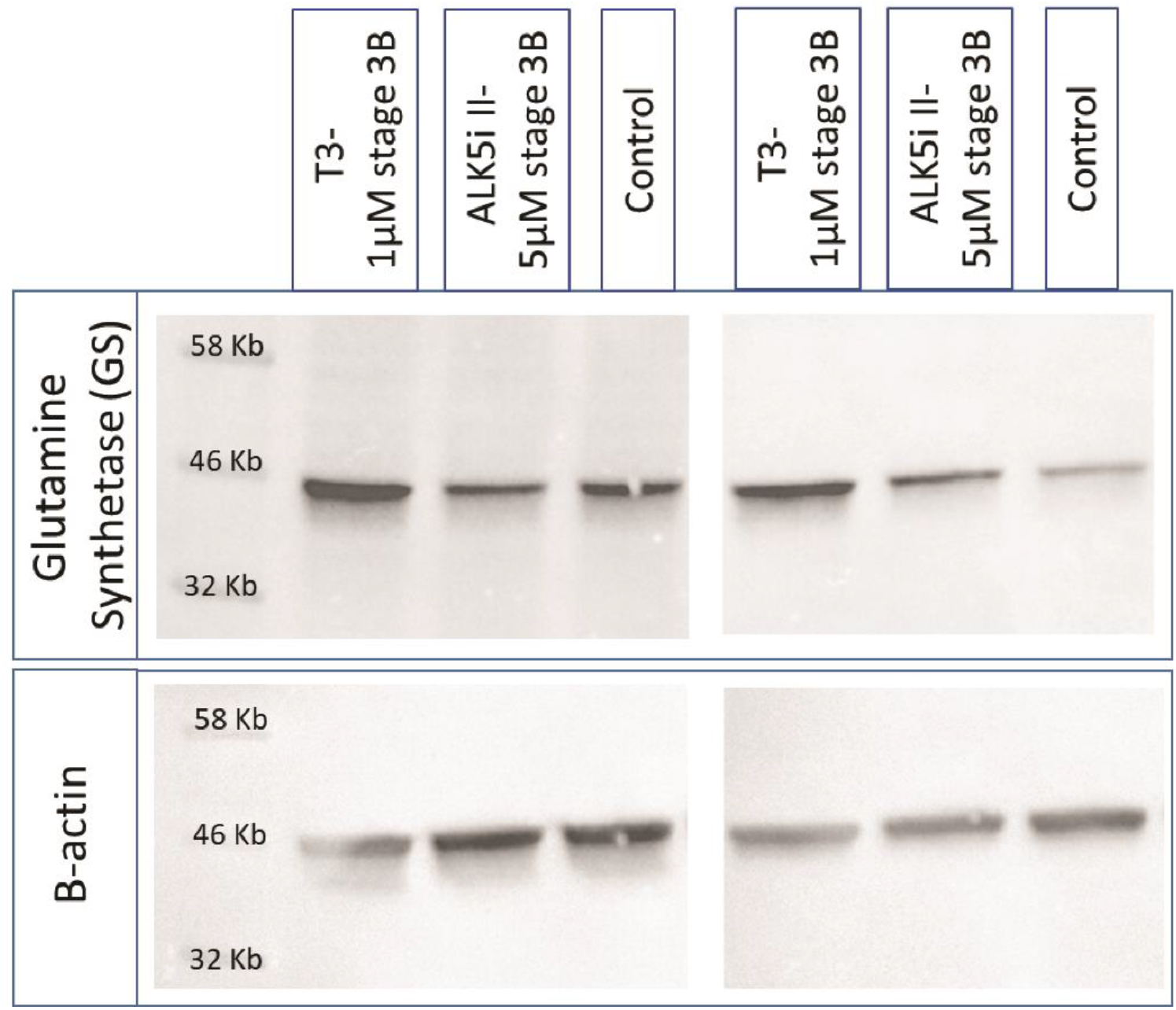
Glutamine synthetase Western Blot. Western Blots against Glutamine synthetase on hepatocyte-like cells treated with 1 µM T3 during Stage 3B, 5 µM ALK5i II during Stage 3B and vehicle control. Antbody against B-actin used as loading control. Data not quantified.

## References

1. Rowe, C., et al., Proteome-wide analyses of human hepatocytes during differentiation and dedifferentiation. Hepatology, 2013. 58(2): p. 799–809.

2. D’Amour, K.A., et al., Efficient differentiation of human embryonic stem cells to definitive endoderm. Nat Biotech, 2005. 23(12): p. 1534–1541.

3. Hay, D.C., et al., Efficient Differentiation of Hepatocytes from Human Embryonic Stem Cells Exhibiting Markers Recapitulating Liver Development In Vivo. STEM CELLS, 2008. 26(4): p. 894–902.

4. Si-Tayeb, K., et al., Highly efficient generation of human hepatocyte–like cells from induced pluripotent stem cells. Hepatology, 2010. 51(1): p. 297–305.

5. Song, Z., et al., Efficient generation of hepatocyte-like cells from human induced pluripotent stem cells. Cell Res, 2009. 19(11): p. 1233–1242.

6. Cai, J., et al., Directed differentiation of human embryonic stem cells into functional hepatic cells. Hepatology, 2007. 45(5): p. 1229–1239.

7. Touboul, T., et al., Generation of functional hepatocytes from human embryonic stem cells under chemically defined conditions that recapitulate liver development. Hepatology, 2010. 51(5): p. 1754–1765.

8. Agarwal, S., K.L. Holton, and R. Lanza, Efficient Differentiation of Functional Hepatocytes from Human Embryonic Stem Cells. STEM CELLS, 2008. 26(5): p. 1117–1127.

9. Basma, H., et al., Differentiation and transplantation of human embryonic stem cell-derived hepatocytes. Gastroenterology, 2009. 136(3): p. 990–9.

10. Chen, Y.-F., et al., Rapid generation of mature hepatocyte-like cells from human induced pluripotent stem cells by an efficient three-step protocol. Hepatology. 55(4): p. 1193–1203.

11. Hannan, N.R., et al., Production of hepatocyte-like cells from human pluripotent stem cells. Nat Protoc, 2013. 8(2): p. 430–7.

12. Avior, Y., et al., Microbial-derived lithocholic acid and vitamin K2 drive the metabolic maturation of pluripotent stem cells–derived and fetal hepatocytes. Hepatology, 2015. 62(1): p. 265–278.

13. Baxter, M., et al., Phenotypic and functional analyses show stem cell-derived hepatocyte-like cells better mimic fetal rather than adult hepatocytes. Journal of Hepatology, 2015. 62(3): p. 581–589.

14. Schwartz, R.E., et al., Pluripotent stem cell-derived hepatocyte-like cells. Biotechnology Advances, 2014. 32(2): p. 504–513.

15. Zhao, D., et al., Promotion of the efficient metabolic maturation of human pluripotent stem cell-derived hepatocytes by correcting specification defects. Cell Res, 2013. 23(1): p. 157–161.

16. Nakamori, D., et al., Hepatic maturation of human iPS cell-derived hepatocyte-like cells by ATF5, c/EBPα, and PROX1 transduction. Biochemical and Biophysical Research Communications, 2016. 469(3): p. 424–429.

17. Godoy, P., et al., Gene networks and transcription factor motifs defining the differentiation of stem cells into hepatocyte-like cells. Journal of Hepatology, 2015. 63(4): p. 934–942.

18. Tauran, Y., et al., Analysis of the transcription factors and their regulatory roles during a step-by-step differentiation of induced pluripotent stem cells into hepatocyte-like cells. Mol Omics, 2019. 15(6): p. 383–398.

19. Jing, R., C.B. Duncan, and S.A. Duncan, A small-molecule screen reveals that HSP90β promotes the conversion of induced pluripotent stem cell-derived endoderm to a hepatic fate and regulates HNF4A turnover. Development, 2017. 144(10): p. 1764–1774.

20. Shan, J., et al., Identification of small molecules for human hepatocyte expansion and iPS differentiation. Nat Chem Biol, 2013. 9(8): p. 514–520.

21. Gerrard, D.T., et al., An integrative transcriptomic atlas of organogenesis in human embryos. eLife, 2016. 5: p. e15657.

22. Du, Y., et al., Human Hepatocytes with Drug Metabolic Function Induced from Fibroblasts by Lineage Reprogramming. Cell Stem Cell, 2014. 14(3): p. 394–403.

23. Trapnell, C., L. Pachter, and S.L. Salzberg, TopHat: discovering splice junctions with RNA-Seq. Bioinformatics (Oxford, England), 2009. 25(9): p. 1105–1111.

24. Harrow, J., et al., GENCODE: the reference human genome annotation for The ENCODE Project. Genome research, 2012. 22(9): p. 1760–1774.

25. Bolstad, B.M., preprocessCore: A collection of pre-processing functions. Bioconductor, 2017.

26. Robinson, M.D., D.J. McCarthy, and G.K. Smyth, edgeR: a Bioconductor package for differential expression analysis of digital gene expression data. Bioinformatics, 2010. 26(1): p. 139–40.

27. Alexa, A.R. J, topGO: Enrichment Analysis for Gene Ontology. R package version 2.26.0. BioConductor, 2016.

28. Nyirenda, M.J., et al., Prenatal programming of hepatocyte nuclear factor 4alpha in the rat: A key mechanism in the ‘foetal origins of hyperglycaemia’? Diabetologia, 2006. 49(6): p. 1412–20.

29. Harries, L.W., et al., The Diabetic Phenotype in HNF4A Mutation Carriers Is Moderated By the Expression of HNF4A Isoforms From the P1 Promoter During Fetal Development. Diabetes, 2008. 57(6): p. 1745–1752.

30. Hashimoto, K., et al., Liver X Receptor-α Gene Expression Is Positively Regulated by Thyroid Hormone. Endocrinology, 2007. 148(10): p. 4667–4675.

31. Hashimoto, K. and M. Mori, Crosstalk of thyroid hormone receptor and liver X receptor in lipid metabolism and beyond [Review]. Endocr J, 2011. 58(11): p. 921–30.

32. Chader, G.J., Hormonal effects on the neural retina: I. Glutamine synthetase development in the retina and liver of the normal and triiodothyronine-treated rat. Archives of Biochemistry and Biophysics, 1971. 144(2): p. 657–662.

33. Gebhardt, R. and D. Mecke, Heterogeneous distribution of glutamine synthetase among rat liver parenchymal cells in situ and in primary culture. Embo j, 1983. 2(4): p. 567–70.

34. Braeuning, A., et al., Differential gene expression in periportal and perivenous mouse hepatocytes. The FEBS Journal, 2006. 273(22): p. 5051–5061.

35. Ishikawa, F., K. Nose, and M. Shibanuma, Downregulation of hepatocyte nuclear factor-4α and its role in regulation of gene expression by TGF-β in mammary epithelial cells. Experimental Cell Research, 2008. 314(10): p. 2131–2140.

36. de Lucas, S., et al., Nitric oxide and TGF-beta1 inhibit HNF-4alpha function in HEPG2 cells. Biochem Biophys Res Commun, 2004. 321(3): p. 688–94.

37. Antoniou, A., et al., Intrahepatic bile ducts develop according to a new mode of tubulogenesis regulated by the transcription factor SOX9. Gastroenterology, 2009. 136(7): p. 2325–2333.

38. Yamasaki, H., et al., Suppression of C/EBPalpha expression in periportal hepatoblasts may stimulate biliary cell differentiation through increased Hnf6 and Hnf1b expression. Development, 2006. 133(21): p. 4233–43.

39. Bogacheva, M.S., M.A. Bystriakova, and Y.-R. Lou, Thyroid Hormone Effect on the Differentiation of Human Induced Pluripotent Stem Cells into Hepatocyte-Like Cells. Pharmaceuticals, 2021. 14(6).

40. Ma, H., et al., The nuclear receptor THRB facilitates differentiation of human PSCs into more mature hepatocytes. Cell Stem Cell, 2022.

41. Menéndez-Hurtado, A., A. Santos, and A. Pérez-Castillo, Characterization of the Promoter Region of the Rat CCAAT/Enhancer-Binding Protein α Gene and Regulation by Thyroid Hormone in Rat Immortalized Brown Adipocytes. Endocrinology, 2000. 141(11): p. 4164–4170.

42. Jurado, L.A., et al., Conserved Amino Acids within CCAAT Enhancer-binding Proteins (C/EBPα and β) Regulate Phosphoenolpyruvate Carboxykinase (PEPCK) Gene Expression. Journal of Biological Chemistry, 2002. 277(31): p. 27606–27612.

43. Menéndez-Hurtado, A., et al., Regulation by Thyroid Hormone and Retinoic Acid of the CCAAT/Enhancer Binding Protein α and β Genes during Liver Development. Biochemical and Biophysical Research Communications, 1997. 234(3): p. 605–610.

44. Sinha, R.A., B.K. Singh, and P.M. Yen, Direct effects of thyroid hormones on hepatic lipid metabolism. Nature reviews. Endocrinology, 2018. 14(5): p. 259–269.

45. Cvoro, A., et al., A thyroid hormone receptor/KLF9 axis in human hepatocytes and pluripotent stem cells. Stem cells (Dayton, Ohio), 2015. 33(2): p. 416–428.

46. Feng, X., et al., Thyroid Hormone Regulation of Hepatic Genes in Vivo Detected by Complementary DNA Microarray. Molecular Endocrinology, 2000. 14(7): p. 947–955.

47. Otto, T. and J. Fandrey, Thyroid Hormone Induces Hypoxia-Inducible Factor 1α Gene Expression through Thyroid Hormone Receptor β/Retinoid X Receptor α-Dependent Activation of Hepatic Leukemia Factor. Endocrinology, 2008. 149(5): p. 2241–2250.

48. Rezania, A., et al., Reversal of diabetes with insulin-producing cells derived in vitro from human pluripotent stem cells. Nat Biotech, 2014. advance online publication.

49. Christoffolete, M.A., et al., Regulation of thyroid hormone activation via the liver X-receptor/retinoid X-receptor pathway. The Journal of endocrinology, 2010. 205(2): p. 179–186.

50. Clotman, F., et al., Control of liver cell fate decision by a gradient of TGF beta signaling modulated by Onecut transcription factors. Genes & development, 2005. 19(16): p. 1849–1854.

